# Brain oscillations track the formation of episodic memories in the real world

**DOI:** 10.1101/042929

**Authors:** Benjamin Griffiths, Ali Mazaheri, Stefan Debener, Simon Hanslmayr

## Abstract

Despite the well-known influence of environmental context on episodic memory, little has been done to enhance contextual richness within the lab. This leaves a blind spot lingering over the neuronal correlates of episodic memory formation in the real world. To address this, we presented participants with series of words to memorise along a pre-designated route across campus. Meanwhile, a mobile EEG system acquired the associated neural activity. Replicating lab-based subsequent memory effects (SMEs), we identified significant low-frequency power decreases, including beta power decreases over the left inferior frontal gyrus. Additionally, the paradigm enabled us to dissociate the oscillatory correlates of temporal and spatial clustering. Specifically, we found spatially clustered items exhibited significantly greater theta power decreases within the left medial temporal lobe than temporally clustered items. These findings go beyond lab-based studies, which are limited in their capabilities to investigate environmental contextual factors that guide memory formation.

## Introduction

Episodic memory refers to rich memories of personally experienced events. The details of these memories not only encompass the event itself but also the environmental setting surrounding the event, such as where and when the event occurred. For example, when recalling the last movie you saw, the act of retrieving this memory may also lead to the incidental recall of the cinema you went to and at what time of day you saw the movie. Unfortunately, the examination of the neural correlates of episodic memory occurs most often within a shielded magnetoencephalogram (MEG) suite or a magnetic resonance imaging (MRI) scanner, a setting in no way representative of the world typically experienced in day-to-day life. As environmental setting has consistently been shown to be a fundamental factor in episodic memory (Baddeley & Godden, 1975; Smith & Vela, 2001), static laboratory settings have impeded a more comprehensive understanding of episodic memory. Indeed, reliance on these lab-based environments means the field has a “blind spot” hovering over episodic memory formation in the real world. While it is impossible to acquire MEG or MRI in daily-life settings, true progress has been made in the use of portable EEG outdoors (De Vos, Gandras, & Debener, 2014; Debener, Minow, Emkes, Gandras, & de Vos, 2012). Embracing these advances, we aimed to identify the electrophysiological correlates of episodic memory formation in the real world.

The most common approach to studying episodic memory formation is the subsequent memory effect (SME). SMEs are the neural signature of successful memory formation, calculated by contrasting the neural activity at encoding which predicts later remembering with the activity that predicts later forgetting, hence isolating the activity unique to memory formation. Oscillatory SMEs are in part characterised by low frequency (<20Hz) power decreases (Burke et al., 2013; Fell, Ludowig, Rosburg, Axmacher, & Elger, 2008; Fellner, Bäuml, & Hanslmayr, 2013; Greenberg, Burke, Haque, Kahana, & Zaghloul, 2015; Guderian, Schott, Richardson-Klavehn, & Duzel, 2009; Klimesch et al., 1996; Long & Kahana, 2015; Noh, Herzmann, Curran, & De Sa, 2014), although this may be an oversimplification of theta band activity during encoding (Nyhus & Curran, 2010; Staudigl & Hanslmayr, 2013). Certainly, beta power (13-20Hz) decreases have been shown to reliably arise over task-relevant sensory regions during successful memory formation, a result that has been attributed to information processing (Hanslmayr, Staudigl, & Fellner, 2012). Critically, a recent EEG-repetitive transcranial magnetic stimulation (rTMS) study has demonstrated that beta power decreases are causally relevant to this process (Hanslmayr, Matuschek, & Fellner, 2014). Given the proposed role of beta in processing task relevant information, unsurprisingly the spatial topography varies depending on stimulus type. For example, verbal materials elicit beta power decreases over the left prefrontal cortex (Hanslmayr, Spitzer, & Bauml, 2009) whereas visual materials elicit slower frequency decreases over parietal and occipital regions (Noh et al., 2014). Nevertheless, the predictability of these beta power decreases following successful memory formation provides a reliable benchmark to contrast with real world recordings in order to identify whether lab-based SME findings are replicable in the real world.

Beyond the replication of previous lab-based findings, portable EEG technology allows the investigation of aspects of episodic memory that only occur in their entirety in the real world, such as the contextual clustering phenomenon. Contextual clustering is the grouped recalling of several events based on a shared contextual detail. Returning to the movie example above, recalling where you saw your last movie in may also lead to the incidental recall of other movies you have seen in the same cinema. Most commonly, contextual clustering has been demonstrated for events which share a similar temporal context (i.e. events that occurred at similar points in time; Howard & Kahana, 2002). However, there is growing evidence that contextual clustering can occur outside the time domain (Polyn, Norman, & Kahana, 2009). Of particular relevance to the experiment presented here, recent studies have demonstrated spatial contextual clustering (i.e. events that occurred at similar points in space; Copara et al., 2014; Miller, Lazarus, Polyn, & Kahana, 2013). To date however, the predominant use of virtual reality in these studies means that the richness of spatial context experienced in real life is ignored. Furthermore, these studies have relied on random travel patterns to dissociate spatial and temporal contextual effects. Unfortunately, a large number of random trajectories would inevitably mean that spatial and temporal context incidental correlate at various points during the experiment, introducing a confounding variable and potentially trivial explanation of spatial clustering. Indeed, it remains unclear to what extent spatial context affects episodic memory in day-to-day life. Here, the use of novel navigational paths that disentangle temporal and spatial context in real-world environments overcomes these pitfalls and aims to further evidence for spatial contextual clustering.

On a neuronal level, contextual clustering occurs (in part) during memory encoding. Long and Kahana (2015) found that later temporal clustering correlates with gamma power increases in the left inferior temporal gyrus and the hippocampus during encoding. However, to the best of our knowledge, no experiment has yet to deduce whether these patterns of activation are unique to subsequent temporal clustering or a part of a more general clustering mechanism. Furthermore, the neural correlates of subsequent spatial contextual clustering remain unknown. Our *a priori* assumptions are that subsequent temporal and spatial clustering would encompass the medial temporal lobe (MTL) – the home of place and time cells (Eichenbaum, 2014; MacDonald, Lepage, Eden, & Eichenbaum, 2011; O’Keefe, 1976). Given the intimate relationship between place cells and theta band activity, it may also be plausible to suggest that the spatial clustering effect would be observable within the theta frequency (Burgess & O’Keefe, 2011; O’Keefe & Recce, 1993).

In this experiment, we address two important questions; 1) do oscillatory lab-based episodic memory studies possess ecological validity? and 2) what are the neural correlates of contextual clustering in real world environments? Following a predefined route and led by the experimenter (see figure 1A and 1B), participants were presented with words to encode and associate with their current location (see figure 1C), a situation similar to remembering several text messages on the way to the supermarket. Participants were shown 4 lists of 20 words, where each list was presented on a spiralling route. These spiralling routes disentangled spatial and temporal context, i.e. as time progressed (temporal context), the participant continually exited and returned to a local environment (spatial context). Glancing at figure 1A, it is apparent that while *‘b’* is closest to ‘*a*’ in terms of temporal proximity (and therefore sharing greater temporal contextual similarity), ‘*e*’ is closest to ‘*a*’ in terms of spatial proximity (and hence is sharing greater spatial contextual similarity). After being shown a list of words, participants were removed from the environment and completed a free recall test. Finally, participants guided the experimenter to where they thought each recalled word was shown and the location was marked by GPS. We aim to replicate the ubiquitous low frequency power decreases in lab-based subsequent memory studies (Hanslmayr & Staudigl, 2014), in particular the beta power decreases over the left inferior frontal gyrus elicited by verbal SME paradigms (Hanslmayr et al., 2011). Furthermore, we aim to identify and dissociate the neural correlates of spatial and temporal contextual encoding. To this end, we conducted a ROI analysis of source activity within the medial temporal lobe, which contrasted neural activity associated with subsequent temporal clustering with that of subsequent spatial clustering. In short, this is the first experiment directly observing the neural correlates of episodic memory encoding in the real world, allowing both the validation of a large body of episodic memory literature and the identification of how real world context affects the neural correlates of encoding.

**Figure 1.**
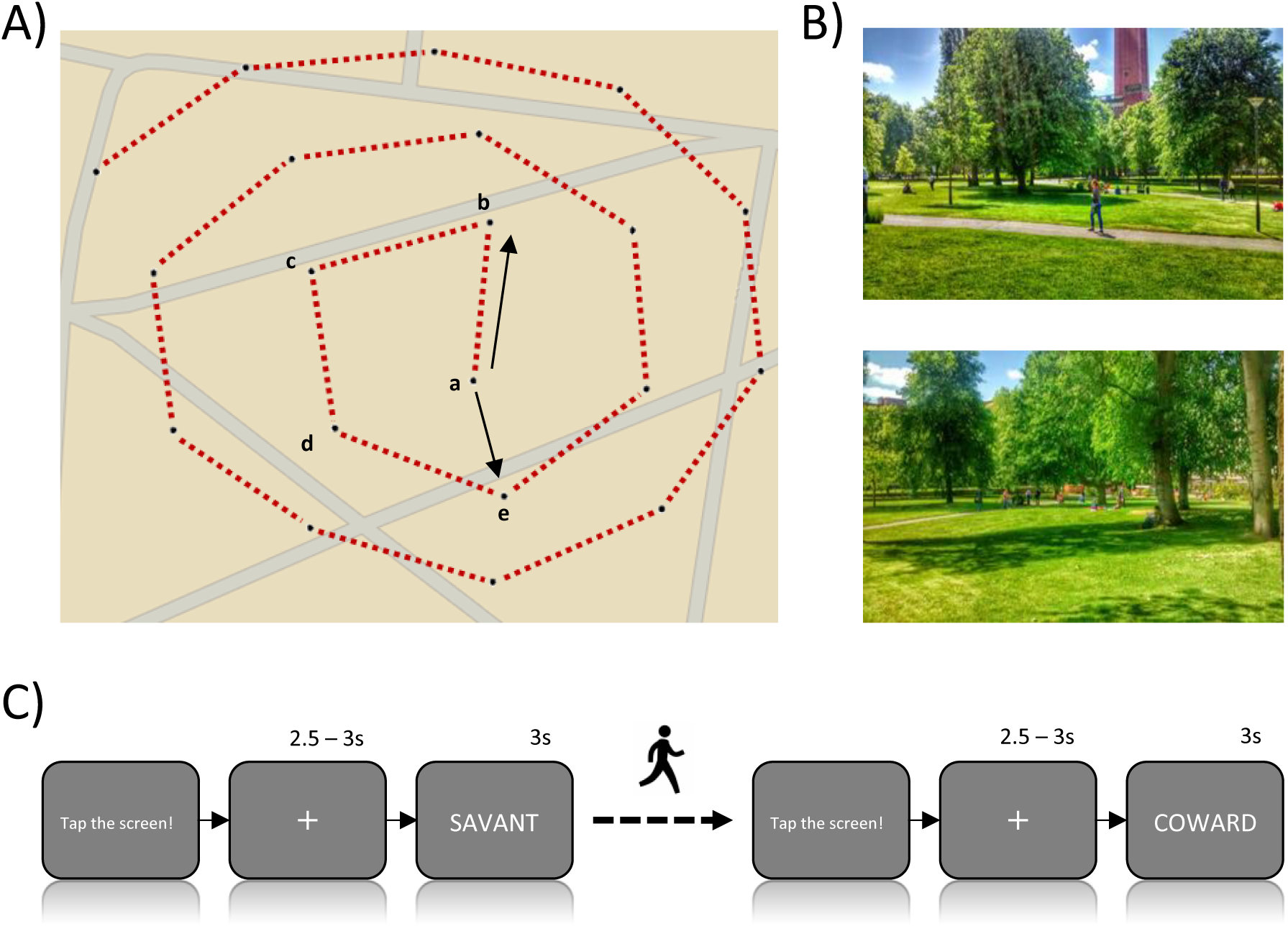
Behavioural paradigm. A) Spiral path. Participants were guided along the red line by the experimenter. At each black dot, the participant was shown one word to encode along with the presentation location. The arrows depict approximate distance between two black dots; the Euclidean distance between the black dots ‘a’ and ‘e’ is less than the two sequential black dots, ‘a’ and ‘b’, dissociating spatial and temporal order. B) Example pictures of the campus areas where the experiment took place. C) A visual representation of each trial as shown on the tablet screen. After the experimenter tapped the screen, a word was displayed following a variable fixation window. Participants were then shown to the next location (black dot in A) and the process was repeated.

## Results

### Behavioural Results

On average, participants recalled 50.50% of each 20 word list and when attempting to locate where each word was presented, were on average 14.74 metres away from the presentation location. To assess spatial and temporal clustering, we devised a novel ‘contextual error’ term. Contextual error describes the observed contextual shift between sequentially recalled items relative to the shortest possible contextual shift, providing a standardised measure that allows direct contrasts between temporal and spatial clustering (see methods for details). The greater the contextual error, the smaller the contextual clustering observed in recall patterns. Eighty percent of participants showed less temporal contextual error (i.e. more temporal contextual clustering) than spatial contextual error (see figure 2). A one-sample t-test revealed significantly greater spatial clustering than expected by chance, t(19)=-4.349, p<0.001, Cohen’s d=1.070, 95% CI [-1.784, ‐.625], matching previous virtual reality results (Miller, Lazarus, et al., 2013). Furthermore, another one-sample t-test revealed significantly greater temporal clustering than expected by chance, t(19)=-5.811, p<0.001, Cohen’s d=1.684, 95% CI [-4.043, ‐1.902], again matching previous lab-based findings (e.g. Long & Kahana, 2015). A dependent samples t-test revealed significantly greater temporal clustering than spatial clustering, t(19)=-4.654, p<0.001, Cohen’s d=0.961, 95% CI [-2.563, ‐.973]. To examine how contextual error relates to memory performance, the mean contextual error of each participant was correlated with their average hit-rate and spatial accuracy. Temporal contextual error did not correlate with memory performance in the free recall task (r=-0.283, p=0.227) or spatial accuracy when returning to presentation locations (r=-0.232, p = 0.326); the same was true for spatial contextual error (free recall performance: r=-0.230, p=0.330; spatial accuracy: r=-0.280, p=0.233).

**Figure 2.**
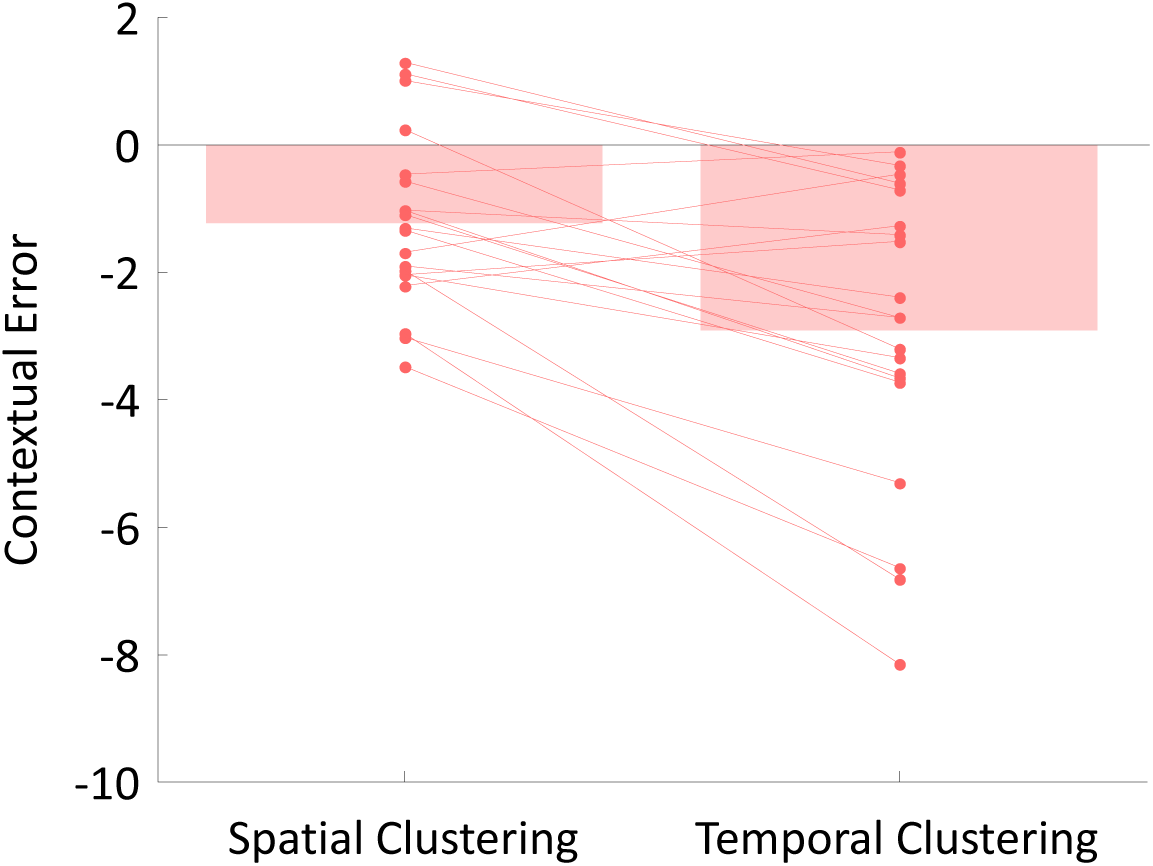
Bar plots represent the mean spatial and temporal ‘contextual error score’. A smaller score represents greater contextual clustering. The amount of contextual clustering expected by chance is zero. Individual scatter points represent the contextual error score of each participant. Spatial and temporal clustering was significantly greater than chance (p<0.001). Temporal was significantly greater than spatial clustering (p<0.001).

### Subsequent Memory Analysis

Given the robust nature of beta power decreases over relevant sensory regions during memory formation, we first aimed to replicate a key lab-based finding in verbal episodic memory studies: a beta power (15-20Hz) decrease over the left inferior frontal gyrus within 1 second of stimulus onset (Hanslmayr et al., 2011). Using a cluster-based permutation test to control for multiple comparisons (see Maris & Oostenveld, 2007), a one-tailed dependent samples t-test revealed a significant power decrease for hits in comparison to misses between 0 and 1 second post stimulus (p=0.009; see figure 3A and 3B). To identify whether this beta power decrease arose in the left inferior frontal gyrus, the 1 second window was then reconstructed on source level, undergoing the same analytical procedure as its sensor level counterpart. A one-tailed dependent samples t-test revealed a significant power decrease for hits in comparison to misses (p=0.026). We determined peak activity by sliding a 6mm radius sphere across the significant cluster and calculating the sum of activity within this sphere (see methods for details); these results were confirmed by visual inspection of the 1% of most extreme voxels within the major cluster. Peak activity was localised to left superior and middle temporal poles, [MNI coord. x=-40, y=19, z=-30; ~BA 38], while visual inspection of the most extreme 1% of voxels within the significant cluster revealed a further difference between later remembered and later forgotten items in the left inferior frontal gyrus (IFG), [MNI coord. x=-39, y=30, z=-18; ~BA 47], (see figure 3C). These results replicate the previous findings of beta power decreases over the left IFG following successful memory formation of verbal information (Hanslmayr et al., 2011, 2009).

**Figure 3.**
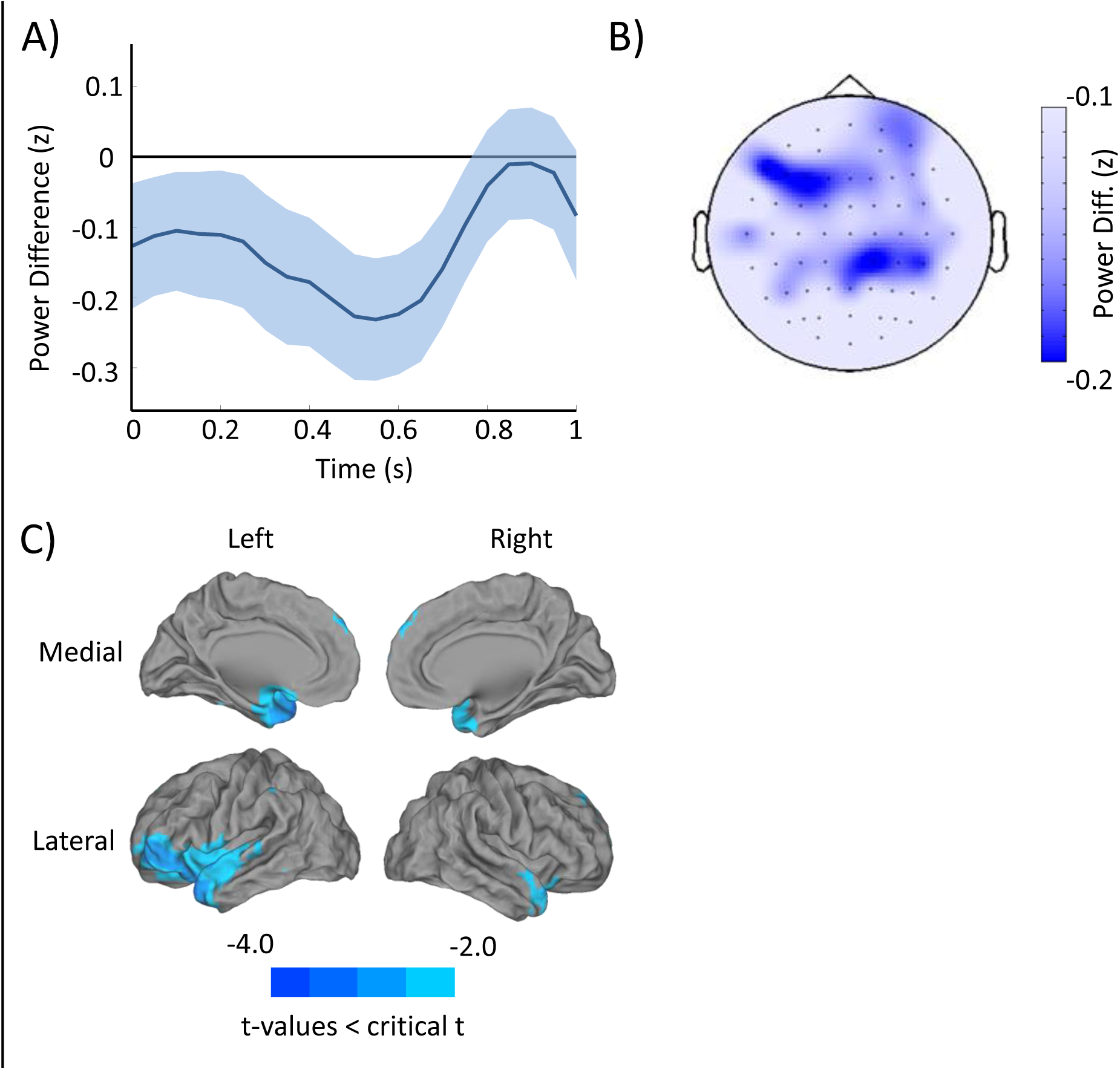
SME (hits minus misses) in the a priori region of interest [0 to 1000ms, 15 to 20Hz]. A) The time course of z-transformed power differences between the later remembered (hits) and later forgotten (misses) items, averaged over channels and frequency bins with standard error of the mean. B) Topography of power differences between hits and misses, averaged across the a priori time-frequency window. C) Source localisation of a priori window of interest. Differences show a significantly greater beta power decrease in the hits condition over left inferior frontal regions.

In order to provide a more comprehensive picture of the low-frequency SMEs in the real world, a sliding window analytical approach was used (Staudigl & Hanslmayr, 2013) where cluster-based permutation tests controlled for multiple comparisons within each window (Maris & Oostenveld, 2007) and false discovery rate (FDR; Benjamini & Hochberg, 1995) controlled for multiple comparisons across windows. Given the ubiquitous nature of power decreases within the alpha and beta bands accompanying successful memory formation (Hanslmayr et al., 2012), one-tailed dependent samples t-tests were used to analysis the subsequent memory effect between 8 and 30 Hz. As some controversy surrounds theta band activity, two-tailed dependent samples t-tests where used for frequencies between 3 and 7 Hz. Analysis revealed significant, FDR corrected, p-values (p_corr_<0.05) across the frequency and time spectrum (see figure 4A). Specifically, low frequency theta (3-4Hz, p_corr_<0.05) power decreases for hits in comparison to misses were observed between 250ms and 950ms post-stimulus; alpha (8-12Hz, p_corr_<0.05) power decreases for hits in comparison to misses were observed between 400ms and 800ms post-stimulus; and beta (21-25Hz) power decreases for hits in comparison to misses were observed just before (-250ms to 0ms, p_corr_<0.05) and later after stimulus onset (1000-1300ms, p_corr_<0.05). These low frequency power decreases match many other effects reported in the literature (see Hanslmayr & Staudigl, 2014). The difference in power between subsequently remembered versus forgotten items did not correlate with spatial accuracy for presentation locations.

**Figure 4.**
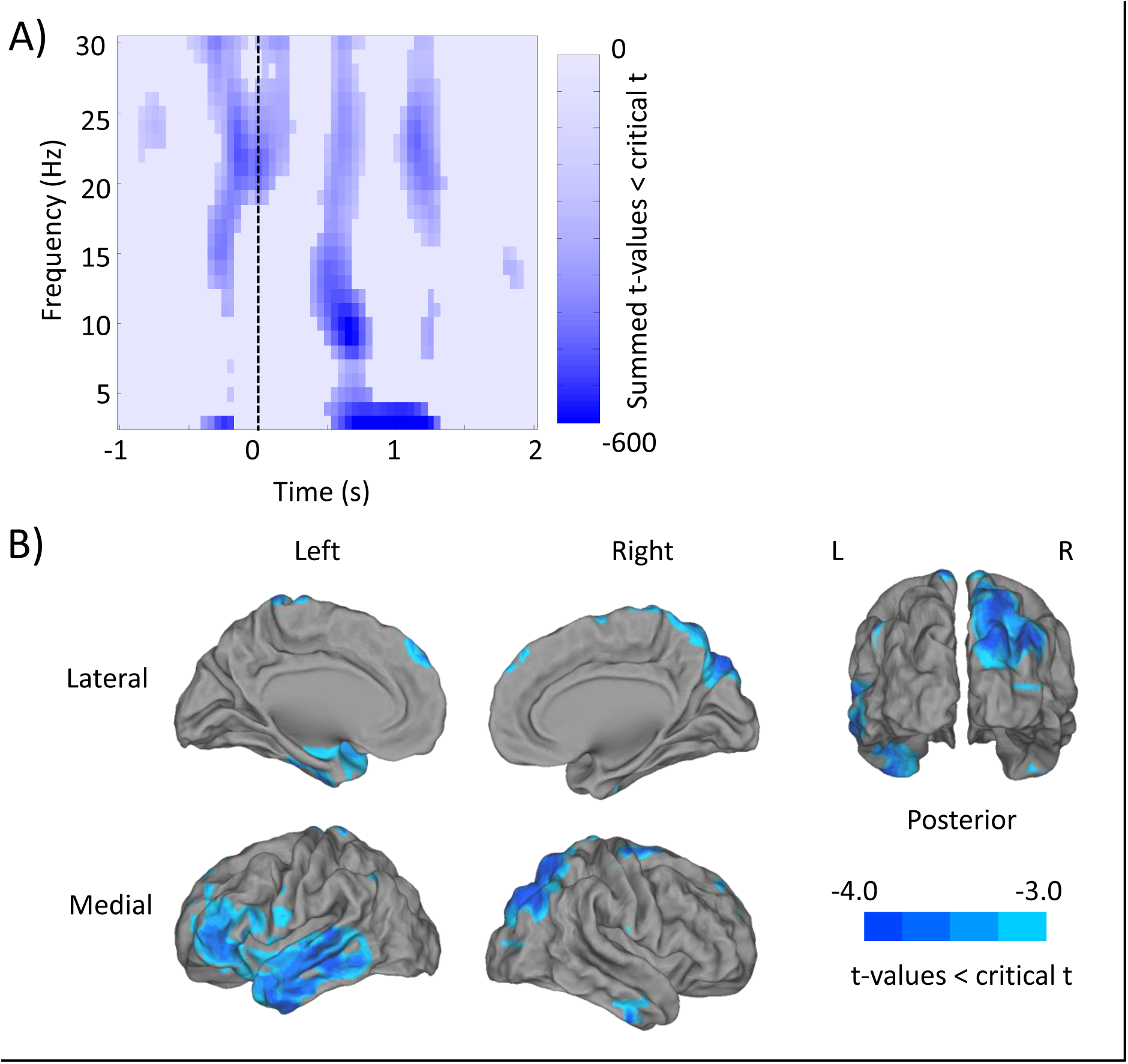
Subsequent memory effect (hits – misses) across the entire data spectrum. A) Time-frequency representation of cluster t-values for each significant sliding window. All non-significant FDR corrected time-frequency windows are masked. B) Source localisation of the significant theta effect (3-4Hz) between 600ms and 1200ms post-stimulus.

Significant regions of activity observed on sensor-level were then reconstructed on source level. Theta activity (3-4Hz, 600-1200ms, p=0.005) peaked in the right superior occipital area, the right precuneus and the right cuneus, [MNI coord. x=19, y=-87, z=39; ~BA 19]. Visual inspection of the theta source activity also revealed peak activity within the left middle and inferior temporal gyri, [MNI coord. x=-52, y=-10, z=-26; ~BA 20], and the right superior parietal lobe, [MNI coord. x=25, y=-64, z=53; ~BA 7] (see figure 4B). Generally speaking, these theta power decreases occurred in regions associated with task-relevant stimuli processing (i.e. semantic processing, Pobric, Lambon Ralph, & Jefferies, 2009; Visser, Jefferies, & Lambon Ralph, 2010; visuospatial processing, Formisano et al., 2002; Sack et al., 2002), conforming to earlier findings (Greenberg et al., 2015). Alpha power decreases (8-12Hz, 500-800ms, p=0.005) peaked in the right inferior frontal gyrus, the right superior and middle temporal poles and the right insula, [MNI coord. x=40, y=19, z=-27; ~BA 38]. Post-stimulus beta activity (21-25Hz, 1000-1300ms, p=0.003) peaked in the left inferior frontal gyrus, left superior temporal pole and gyrus, and the left rolandic operculum, [MNI coord. x=-58, y=8, z=0; ~BA 48]. Pre-stimulus beta activity (21-25Hz, ‐250-200ms, p=0.003) could not be effectively localised using the spherical cluster, but visual search of the source revealed notable differences in the right superior parietal lobe and right postcentral gyrus, [MNI coord. x=27, ‐50, 58; ~BA 7]. In summary, the real world SME observed here appears to match what is regularly reported in lab-based studies (e.g. Greenberg et al., 2015; Hanslmayr et al., 2009).

### Subsequent Clustering Analysis

Our subsequent clustering analysis was conducted on a time-frequency representation of r-values obtained from correlating power across trials for each channel-frequency-time data point. As a first step, we examined whether the correlation between power and temporal/spatial clustering differed significantly from the null hypothesis (i.e. no correlation; r = 0). Concerning temporal clustering, the sensor level analysis (conducted as in *Subsequent Memory Analysis)* of temporal clustering revealed no cluster exceeding the significance threshold. This is consistent with a previous study which also found no correlation between temporal clustering and low frequency power (Long & Kahana, 2015). Concerning spatial clustering however, a sliding window analysis revealed a cluster consisting of extended slow theta power decreases in the peri-stimulus interval (3-4Hz; ‐1000-1000ms, p_corr_<0.05), and a broader theta post-stimulus power decrease (3-6Hz; 400-900ms, p_corr_<0.05), which predicted spatial clustering (see figure 5A). As above, these windows were reconstructed in source space. The post-stimulus theta activity (3-6Hz; 400-900ms, p_corr_<0.05) peaked in the left calcarine sulcus, cuneus and superior occipital regions, [MNI coord. x=-8, y=-97, z=20; ~BA 17] (see figure 5B). Meanwhile, the peri-stimulus theta activity (3-4Hz; ‐1000-1000ms, p_corr_<0.05), peaked in left superior and medial frontal gyrus, [MNI coord. x=-8, y=39, z=51; ~BA 8] (not pictured due to strong similarity with fig. 5B).

**Figure 5.**
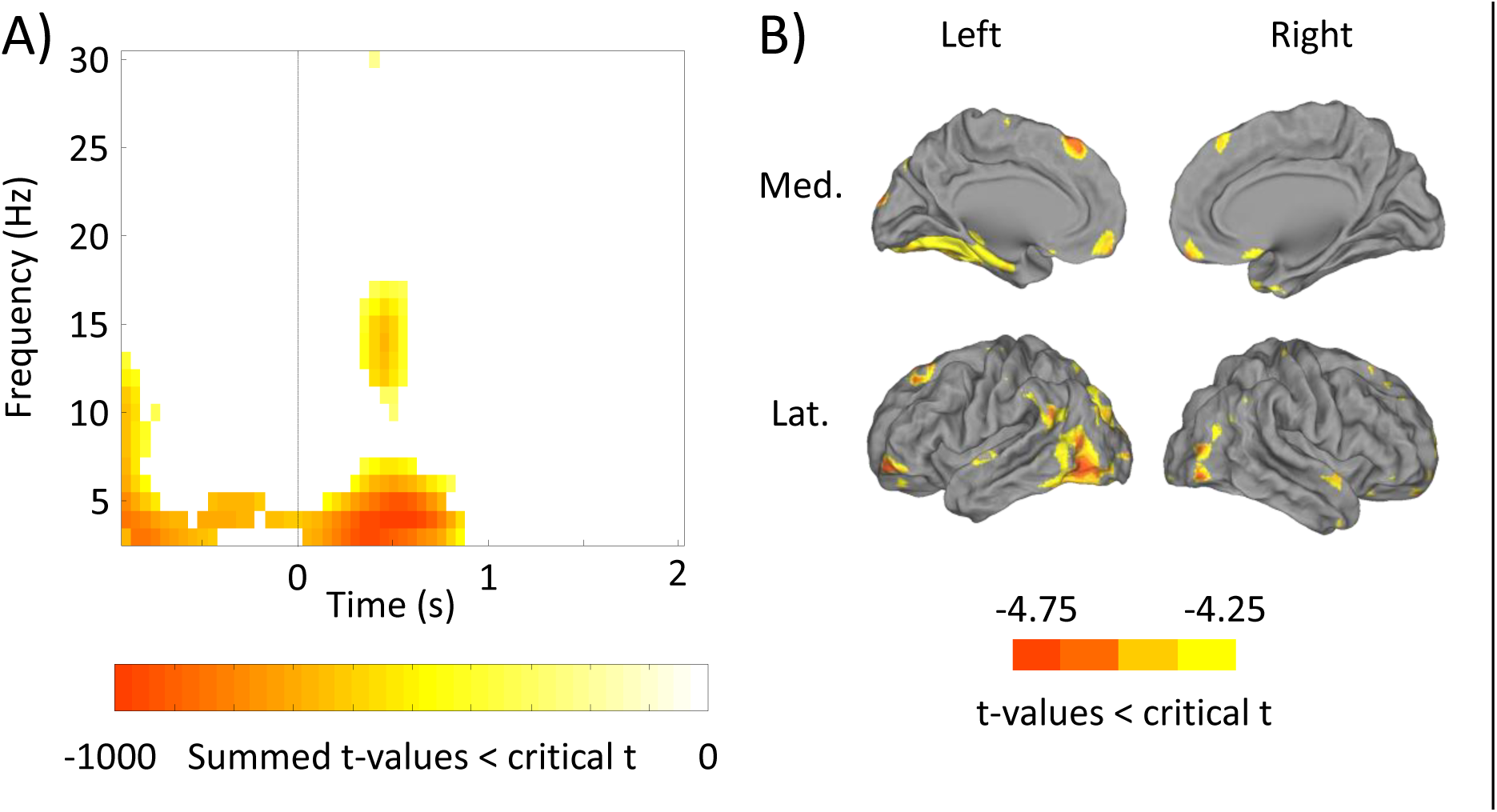
Neural correlates of spatial clustering. A) Time-frequency representation of cluster summed t-values for windows where the observed correlation coefficient was significantly different from the null hypothesis (i.e. r = 0). B) Source reconstruction of the post-stimulus theta power decrease associated with greater spatial clustering (3 – 6Hz, 400 – 900ms).

In a second step, we contrasted the r-values obtained by correlating theta power and temporal clustering with r-values obtained by correlating theta power and spatial clustering, in order to identify whether these theta power decreases were unique to the spatial clustering condition. Cluster analysis indicated that there was a small but significant difference between temporal clustering - theta power effects and spatial clustering - theta power effects (p_corr_<0.05; see figure 6A). T-values indicate that theta power decreases correlate more strongly with spatial clustering than with temporal clustering. When reconstructing this difference on source level (see figure 6B), the spatial-temporal contrast (3-5Hz, 400-800ms, p=0.025) appeared to peak in left frontal superior and medial gyri, [MNI coord. x=-5, y=40, z=57; ~BA 8]. Visual inspection of the peak 1% of activity also revealed greater theta power decreases from within the left medial temporal lobe, [MNI coord. x=-27, y=-1, y=-30; ~BA 36].

**Figure 6.**
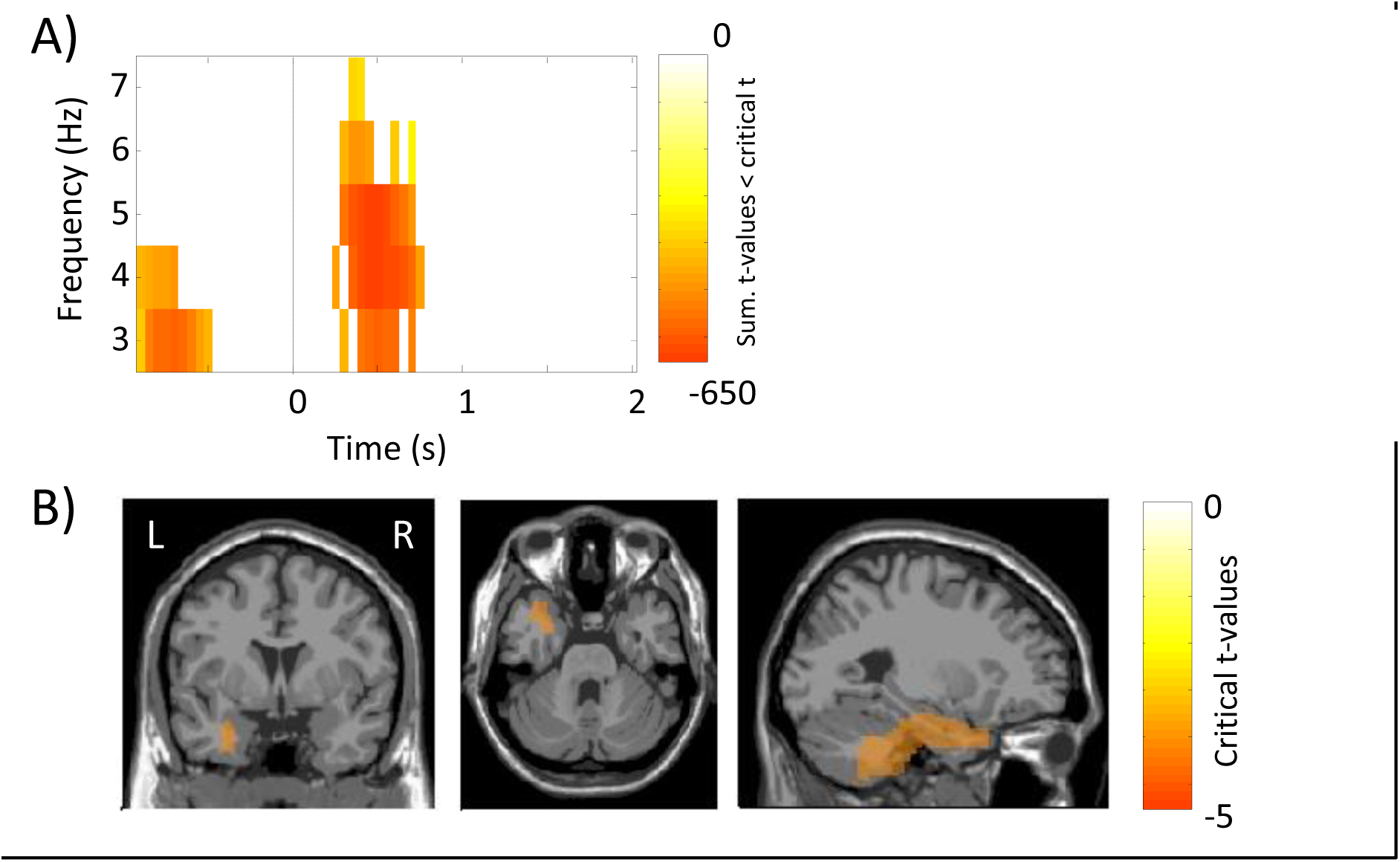
Significant decreases in theta power activity for spatial clustering in comparison to temporal clustering. A) Sensor level time-frequency representation of significant differences in theta power. B) Orthographic plot of source activity differences between spatial clustering and temporal clustering, (left frontal activation occurred on a more medial slice).

## Discussion

This experiment aimed to replicate the oscillatory subsequent memory effect in a real-world environment. Moreover, we investigated the influence of real world contextual factors (i.e. space) on episodic memory relative to contextual factors available within the lab (i.e. time). Participants donned a portable EEG setup and were presented with verbal stimuli on a tablet across the university campus. Each list was presented on a spiral path that disentangled temporal and spatial context. Sorting EEG recordings obtained during encoding by later memory performance allowed the activity unique to successful memory formation to be identified. These results indicated strong beta power decreases over left frontal regions for items which were later remembered in comparison to those which were later forgotten. Furthermore, a broad theta power decrease was observed shortly after stimulus presentation for items later remembered over regions including the left inferior frontal regions and right posterior parietal cortex. Similarly, theta power decreases accompanied strong spatial clustering within the medial temporal lobe in contrast to temporal clustering.

Generally speaking, our real world findings corroborate what others have demonstrated within a lab setting. On a behavioural level, individuals continue to demonstrate both temporal and spatial contextual clustering in an environment where spatial details are significantly richer (Miller, Lazarus, et al., 2013; Miller, Neufang, et al., 2013). Expanding on previous experiments, the spiralling presentation pattern used in this experiment ensured that temporal and spatial context did not overlap. Knowing that temporal clustering could not inform spatial clustering and vice versa, this experiment furthers the notion that temporal clustering and spatial clustering are autonomous.

On an electrophysiological level, we replicated the ubiquitous low-frequency power decreases observed during successful memory formation. Source localisation of the beta power activity revealed decreases in the left frontal and temporal pole regions, both of which are associated with verbal and semantic processing (Pobric et al., 2009). Following the information-via-desynchronisation hypothesis (Hanslmayr et al., 2012), these beta power decreases would reflect verbal information processing necessary for successful memory formation. Although discussed in previous studies (Hanslmayr et al., 2009), given the real world aspects of this study we reiterate that these power decreases (particularly within the beta band) are not viewed as oscillatory correlates of motor activity (Salenius & Hari, 2003). The experimenter ensured the participant was stationary before the presentation of each stimulus, so no motor component should have arisen. If such a component did arise, then it would be evenly distributed between later remembered and later forgotten items, and hence cancel out in the later remembered-later forgotten contrast.

We also observed significant theta power decreases following successful memory formation, particularly for items that demonstrated strong spatial clustering at recall. These power decreases may reflect a common process – possible selective communication within and across spatially diverse regions. Diversity in phase is optimal for communication as signals can arrive at a time of peak excitability and selectively communicate with receiving, down-stream, neural assemblies (Maris, Fries, & van Ede, 2016). There is a wealth of evidence to suggest theta is well suited for such communication needs (see Colgin, 2013 for review). Critically, the diversity in theta phase beneficial for communication would be reflected by theta power decreases in regions relevant to successful memory formation, especially in macro-scopic recording techniques such as EEG. In the context of the current experiment, observed theta power decreases in the temporal poles, posterior parietal cortex and medial temporal regions likely reflect the activation of, and communication between, areas responsible for the processing of semantics (e.g. Whitney, Kirk, O’Sullivan, Lambon Ralph, & Jefferies, 2011), egocentric space (Ciaramelli, Rosenbaum, Solcz, Levine, & Moscovitch, 2010) and allocentric space (Miller, Vedder, Law, & Smith, 2014). Ultimately, these oscillatory dynamics allows the formation of coherent memory episodes. This account would also explain the absence of a similar theta power decrease for temporally clustered items. Temporal clustering might rely on a smaller network involving no communication with spatial processing regions. Consistent with this assumption, a previous study linked temporal clustering to high frequency (gamma) activity which might reflect the action of more local networks (Long & Kahana, 2015). Therefore, reduced activation during temporal clustering would elicit minimal theta power decreases in comparison to those observed following spatial clustering.

In conclusion, our findings are the first to provide strong evidence for the ecological validity of lab-based experiments investigating episodic memory formation and oscillations. More importantly, our investigation into contextual clustering also highlights the significance of real world memory research. We suggest that similar virtual reality studies would not observe such a strong effect of spatial contextual clustering, given the limited spatial information (and spatial congruency) available in virtual reality. Our mobile EEG approach can pave the way towards new insights into the underpinnings of spatial contextual details in newly formed memories that lab-based experiments never could.

## Methods

### Participants

29 students (18-39 years, 69% female) from the University of Birmingham were recruited through a participant pool and rewarded with financial compensation for participation (£25 for a 4 hour session). Nine participants were excluded from the sample due to issues in recording (n=4), poor weather conditions (n=2) or extreme performance in the task (<15 remembered items or <15 forgotten items; n=3). All participants were native English speakers or had lived in an English speaking country for the past 5 years. Participants reported normal or corrected-to-normal vision. Our sample size boundary (n=20) was determined by similar studies which have used samples of ~20 participants to produce reliable oscillatory subsequent memory effects (e.g. Hanslmayr, Spitzer, & Bauml, 2009). A power analysis on pilot behavioural data indicated that a sample size of 16 participants was adequate for detecting a significant behavioural effect using this paradigm (α=0.05; 1– β=0.80). Ethical approval was granted by the University of Birmingham Research Ethics Committee, complying with the Declaration of Helsinki.

### Materials

80 unique abstract nouns and 80 unique locations were split into 4 blocks (20 words and locations per block). The nouns were selected from the MRC Psycholinguistic Database based on scores of low imaginability and concreteness (http://www.psych.rl.ac.uk/). All locations within a block were found in the same large, open space on the university campus. Lists and locations were counterbalanced across participants. Words were presented in black on a light grey background using the OpenSesame experiment builder (2.9.4; Mathôt, Schreij, & Theeuwes, 2012) on a Google Nexus 7 (2013; Google, Mountain View, California) tablet running Android OS (5.1.1). Tones were elicited by the tablet and passed onto a StimTracker (Cedrus Corporation, San Pedro, California), which in turn passed a trigger to the EEG acquisition system. Within each block, the navigated route formed a spiral (although participants were unaware of this; see Figure 1A). Words adjacent in serial order were 20m apart while parallel spiral arms were approximately 15m apart, ensuring words on parallel paths were closer in Euclidean distance than those on the same path. A Garmin eTrex 30 Outdoor Handheld GPS Unit (Garmin Ltd., Canton of Schaffhausen, Switzerland) was used to navigate the route and to mark co-ordinates during the spatial memory test. The GPS could accurately pinpoint a current location to approximately within 3 metres.

### Procedure

Prior to commencing the experiment, participants were informed of the experimental procedure, completed a screening questionnaire and provided informed consent. During the encoding stage of each block, the experimenter guided the participant along a spiral path and at predefined locations presented the participant with a word on the tablet screen. After haptic input from the experimenter (given once the participant was stationary), a fixation cross was displayed in the centre of the screen for 2.5 to 3 seconds (uniformly random), followed by a target word presented for 3 seconds. The lengthy pre-stimulus interval ensured that any motor/motor-rebound effects would not contaminate EEG recordings during the critical presentation window. The participant then encoded the word and the location. After 20 locations had been visited, the participant completed a short subtraction distractor task (“starting at x, count down in steps of y, all the way to zero”) to disrupt any working memory effects. The participant was then guided to a testing cubicle and given 3 minutes to freely recall as many of the words presented as possible. Subsequently, the experimenter led the participant back outside and, using the list of recalled words as a cue, the participant attempted to return to where each word was presented. GPS co-ordinates for each of these recalled locations were recorded. After the participant had recalled as many of the locations as they could remember, the experimenter guided them to the next area to start the following block.

### Behavioural Analysis

Spatial accuracy of recalled locations was determined by calculating the distance between the presentation and recalled locations of each word using the Haversine formula, providing a parametric measure of accuracy in metres. An error term was used to identify whether participants recalled in spatial and/or temporal clusters. ‘Contextual error’ is a novel term which describes the extent to which an individual deviated from the immediate context when recalling events; the smaller the contextual error, the greater the contextual clustering. This provided a standardised, and therefore comparable, measure of temporal and spatial contextual clustering. Contextual error was derived using the equation below:

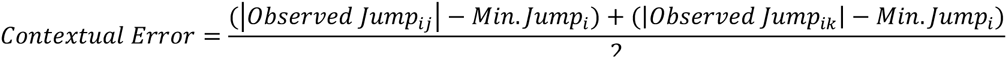

Here, *i* refers to the recalled item under observation, *j* to the item recalled immediately before *i*, and *k* to the item recalled immediately before *j*. *Observed Jump*_*ij*_ and *Observed Jump*_*ik*_ refer to the absolute of the contextual distance between items *i* and *j*, and *i* and *k*, respectively. For example, if items *i* and *j* where presented 30 metres apart, the observed spatial contextual jump at recall would be 30 metres. Spatial distances were measured in metres, while temporal distances were measured by the serial lag between items. As each item within a block was presented approximately 15 seconds after its prior, serial lag and temporal lag are viewed as synonymous. *Min. Jump*_*i*_ refers to the distance between item *i* and the most proximal list item. For example, if the closest item presented to item *i* in space was 12 metres away, the minimum spatial contextual jump would be 12 metres. In the case of temporal context, the minimum jump is always 1 as the jump would be to the immediately preceding/proceeding item presented during encoding. Keeping the example values assigned above, the contextual error (observed jump minus minimum jump) between items *i* and *j* is 18 metres. To compare the two modalities of context, raw contextual error scores were z-transformed using the means and standard deviations of noise data. This noise data was generated by taking the observed hits, randomly assigning a recall position to them, and then calculating the contextual error based on these random recall positions. To provide a measure of clustering rather than idiosyncratic jumps between individual items, an average jump was calculated using the two previously recalled items (hence *ij* and *ik)*. This method is not expected to fundamentally change the results of previous lab-based studies. Lohnas & Kahana (2014) have demonstrated that temporal clustering in free recall is influenced by multiple recent recall items, not the immediately preceding item alone. One-sample t-tests were used to examine whether participants recalled in clusters more greatly than expected by chance. A dependent-samples t-test then compared temporal and spatial contextual error scores.

### EEG Acquisition, Pre-processing and Time-Frequency Decomposition

EEG was recorded using a portable ‘EEGo Sports’ EEG system (ANT Neuro, Enschede, Netherlands) with 65 Ag/AgCl electrodes arranged in a 10/10 system layout (including left and right mastoids, CPz as reference and AFz as ground). Impedances were kept below 20 kΩ, and the sampling rate was 500Hz. For source analysis, head coordinates of all electrodes and the nasion, left pre-auricular area (LPA) and right pre-auricular area of each participant were taken using a Polhemus Fasttrack system (Polhemus, Colchester, VT). The data was preprocessed using the FieldTrip toolbox (Oostenveld, Fries, Maris, & Schoffelen, 2011). The continuous data were epoched into single trials beginning 2000ms before word presentation and ending 3000ms after word presentation. Eye-blinks, saccades and any other consistent artefacts were removed using independent component analysis. Residual irregular artefacts were removed by rejecting the corresponding trials; mean number of trials rejected = 15.45; mean number of hits remaining = 35.25 (max: 51, min. 22); mean number of misses remaining; 26.30 (max. = 42; min. = 16). Bad channels were interpolated based on the data of neighbouring electrodes and the data was given an average reference (mean interpolated = 0.6; max. = 5; min. = 0).

Time-frequency analysis was conducted on the pre-processed dataset for each participant using 7 cycle Morlet wavelets for frequencies of 1 to 30Hz in 1hz steps. The time resolution was set to 50ms. For each frequency-channel pair, the data were z-transformed by first obtaining the average power over time for each trial, and then getting the average and standard deviation of this time-averaged power across trials. This twice-averaged power was then subtracted from the observed power at each channel-frequency pair, and the output was divided by the standard deviation of the time-averaged power. Gaussian smoothing was then applied to the time-frequency representation.

### Subsequent Memory Analysis

Trials were split into two categories; item-location pairs that were later remembered (hits) and later forgotten (misses). The data was first restricted to 0-1000ms post-stimulus between 15 and 20Hz to replicate previous beta power decreases seen in subsequent memory paradigms. Hits and misses for this time-frequency window were contrasted using a dependent samples t-test. A Monte-Carlo randomisation procedure using 2000 permutations was employed to correct for multiple comparisons (see Maris & Oostenveld, 2007). The clusters used in this randomisation procedure were defined by summing the t-values of individual channel-frequency-time triplets that exceeded threshold (α = 0.05).

Subsequently, further power changes in the time-frequency representation were examined. Following previous literature, alpha and beta power decreases were tested, while undirected theta power differences were tested. Accordingly, alpha and beta tests were one-tailed, while theta power tests were two-tailed‥ A sliding window (200ms by 1Hz in size) was passed over the time-frequency window (-1000 to 2000ms), contrasting power differences between hits and misses within the window. The window shifted along the time dimension in steps of 50ms. In this technique, the Monte-Carlo randomisation procedure alone is not sufficient to control for multiple comparisons so the p-values for each sliding window were pooled together and thresholded using false discovery rate (FDR; Benjamini & Hochberg, 1995).

### Subsequent Clustering Analysis

To assess the oscillatory correlates of temporal and spatial clustering during encoding, contextual error scores were correlated with the time-frequency power spectrum. For each participant and for each time-frequency-channel point, the contextual error score for each trial was correlated with the observed power for that trial using a Spearman’s Rank procedure. As less contextual error denotes greater contextual clustering, a negative r-value would indicate a power increase accompanying greater contextual clustering. To aid comprehension, each returned r-value underwent a switching of sign (+0.5 became ‐0.5; ‐0.5 became +0.5), meaning a positive rvalue indicated a power increase with greater contextual clustering. The time-frequency representation of r-values was tested against the null hypothesis that there would be no correlation between power and contextual clustering. This null hypothesis was realised by creating a ‘null data structure’ with the same dimensions as the observed data, but with all observed data points substituted with zeros (i.e. no correlation). The observed data was then contrasted with the ‘null data’ in the same manner as the sliding window approach described above.

### Source Analysis

Observed effects on sensor level were reconstructed in source space using individual head models in combination with the standard MRI and boundary element model (BEM) provided in the FieldTrip toolbox. The Linearly Constrained Minimum Variance (LCMV) beamformer was used to localise sources of significant activity (van Veen, van Drongelen, Yuchtman, & Suzuki, 1997). Pre-processed data was time-locked and then shifted to source space. This placed the time-locked data onto virtual electrodes, which then underwent an identical analytical procedure to its sensor-level counterparts. P-values are presented with each source reconstruction for completeness, but as the time-frequency windows were selected because they exceeded the significance threshold on sensor level, caution should be taken when interpreting source-level p-values. Peak activity was first deduced by sliding a sphere with a 6mm radius over all voxels within the interpolated significant cluster (interpolated grid size: 181x217x181mm). All voxels that fell within the sphere were summed, and the group of voxels that produced the absolute max sum was selected as the peak region. As this approach cannot effectively handle sparse regions of activity, a follow-up visual inspection was conducted. For visual inspection, only the 1% of voxels with the most extreme t-values was examined. The results of visual inspection are only reported when they produced notable differences to the peak sphere approach.

### Additional Analyses

Several further analyses were conducted but were subject to a number of analytical issues. For transparency, these analyses are listed here, but to avoid misinterpretation of the outcomes of these analyses by those glancing over the paper, these results are not reported in the results section. Theta phase to gamma amplitude coupling was investigated using method described by Jiang, Bahramisharif, van Gerven, and Jensen (2015) in an attempt to find similar cross-frequency coupling contextual effects to those reported by Staudigl & Hanslmayr (2013). However, no differences were found, possibly due to the overly noisy gamma activity. Furthermore, differences in source-level connectivity between the medial temporal lobe and the prefrontal cortex for high versus low contextually clustered items was investigated to test the hypothesised neural context model put forward by Polyn & Kahana (2008). Unfortunately, difference in phase angles between virtual electrode connections were almost solely clustered around 0 and π, suggesting a volume conduction issue (Cohen, 2015).

